# Invasive adult jumping worms in Atlantic Canada are chill-susceptible

**DOI:** 10.1101/2024.05.02.592186

**Authors:** Victoria E. Adams, Subash Raj Chettiar, Tanner M. Clow, Emily Gendron, Amber L. Gough, Brianna E. M. Stewart, Erin K. Cameron, Jantina Toxopeus

## Abstract

The jumping worm *Amynthas tokioensis* is invasive in North America, and it has been expanding its range northward in recent years. Because low temperatures typically restrict the geographic distribution of organisms, our goal was to characterize the cold tolerance physiology of adult jumping worms from a site in New Brunswick, Canada (c. 45°N), with the intent of better understanding their geographic range limits. Most of our experiments supported the conclusion that these worms are chill-susceptible: they die during or after exposure to relatively mild low temperatures. When gradually cooled, adult worms lost neuromuscular coordination at approximately 0 °C and froze at a mean temperature of −4.5 °C. They did not survive freezing and showed poor survival following 1 h exposures to 0 °C and subzero temperatures. At higher mild temperatures (5 °C), the worms could survive short (up to 6 h) but not long (e.g., 48 h) chilling durations. We attempted to induce improved cold tolerance via a five-week gradual acclimation to fall-like temperatures, but fall-acclimated worms showed poor survival during and after this acclimation. Acclimation also did not induce accumulation of glucose, a typical cryoprotectant in earthworms. We suggest that *A. tokioensis* can likely persist wherever the growing season is sufficiently warm and long enough for the adults to mature, reproduce, and lay cocoons prior to the chilling temperatures associated with early fall. Future work examining the cold tolerance of the overwintering cocoons will be important for fully understanding the northern range limits of these jumping worms.

## Introduction

*Amynthas tokioensis* (jumping worm) is a pheretimoid earthworm (Oligochaeta: Megascolecidae) native to East and Southeast Asia and invasive in North America (Chang et al. 2021; McAlpine et al. 2022). In their non-native habitats, *A. tokioensis* rapidly alter soil structure and nutrient availability (Chang et al. 2021). *Amynthas* spp. were first reported in Canada in Ontario in 2014 (c. 42°N; Reynolds 2014, 2018), and *A. tokioensis* has since invaded Québec (c. 45°N; Moore and Reynolds 2024) and New Brunswick, Canada (c. 45°N; McAlpine et al. 2022). This species often co-occurs with two other invasive jumping worms, *Amynthas agrestis* and *Metaphire hilgendorfi* (Chang et al. 2018). With their increasingly northward expansion and substantial impacts on soil and other organisms, there is a need to increase our understanding of the thermal limits of these jumping worms.

The thermal limits of jumping worms have been partially described, especially in relation to their annual life cycle. *Amynthas* spp. overwinter as cocoons, hatch as juvenile worms in the early spring, and mature into adults that reproduce and lay cocoons before dying in the fall (Chang et al. 2021). Laboratory experiments have shown that cocoons of *A. agrestis* from Tennessee, USA (c. 35°N) will hatch at 10 °C but not 5 °C (Blackmon et al. 2019). Juvenile *Amynthas* spp. have hatched even after cocoons experience extreme low winter temperatures, *e*.*g*., below −20 °C in Vermont, USA (c. 44°N) during the 2014 polar vortex (Görres et al. 2016). Adult *A. agrestis* do not survive prolonged (28 d) exposures to 5 °C (Tennessee population; Richardson et al. 2009), and the abundance of adult *A. agrestis* in the field declines as air temperatures drop below 10°C, with rapid decline below 5°C (Vermont population; Görres et al. 2014, 2016). While some data on temperature limits have been collected for *A. agrestis*, there is little literature on similar thresholds for the co-invasive *A. tokioensis*, and no information on the thermal limits of Canadian populations.

The lower thermal limits of ectothermic animals can be measured in multiple ways. As environmental temperatures decrease, animals will reach a critical thermal minimum (CTmin), the low temperature at which voluntary movement ceases (Overgaard and MacMillan 2017). Exposure to the CTmin is usually non-lethal (unless for prolonged periods) and lower CTmin values are associated with increased cold tolerance (Overgaard and MacMillan 2017). As environmental temperatures decrease below the CTmin, internal ice formation begins at the supercooling point (SCP; Sinclair et al. 2015), although some species remain active until they freeze (CTmin = SCP; *e*.*g*., Toxopeus et al. 2016). In addition, internal freezing may be initiated by external factors, such as contact with environmental ice (Holmstrup 2003; Toxopeus and Sinclair 2018). Both CTmin and SCP can vary seasonally as animals acclimatize to changing environmental conditions, and in response to acclimation regimes in laboratories (Havird et al. 2020). For example, adult *Eisenia nordenskioldi* (Siberian earthworms) decrease their SCP from −1.2 °C to −3.5 °C both naturally during the fall and when exposed to gradually decreasing temperatures in lab (Meshcheryakova and Berman 2014). The relevance of the SCP for cold tolerance depends on the animal’s cold tolerance strategy.

The two major cold tolerance strategies of ectothermic animals are freeze tolerance and freeze avoidance. Freeze-tolerant animals survive internal ice formation (Holmstrup and Zachariassen 1996; Toxopeus and Sinclair 2018) and temperatures below their SCP, exhibited by adults of the earthworms *E. nordenskioldi* (Holmstrup et al. 1999) and *Dendrobaena octaedra* (Rasmussen and Holmstrup 2002). Freeze-tolerant earthworms often accumulate high levels of glucose as a cryoprotectant molecule (Holmstrup et al. 1999; Slotsbo et al. 2008). Freeze-avoidant (also called freeze-intolerant) animals physiologically suppress the SCP via accumulation of cryoprotectants and antifreeze proteins, but will die if ice forms (i.e., at the SCP), as seen in many terrestrial insects (Toxopeus and Sinclair 2018). Cryoprotective dehydration is an extreme version of freeze avoidance, in which organisms avoid ice formation by substantially decreasing their water content, a strategy used by cocoons of some earthworms (Holmstrup and Westh 1994). Animals that are not cold-tolerant are usually described as chill-susceptible, dying at mild low temperatures above the SCP, such as the earthworms *Eisenia fetida* (red wrigglers; Meshcheryakova and Berman 2014). Chill-susceptible animals may behaviourally avoid low temperatures, for example by overwintering below the frost layer, as seen in the earthworm *Lumbricus terrestris* (nightcrawlers; Nuutinen and Butt 2009). Measuring survival under both short (*e*.*g*., 1 h) and prolonged (*e*.*g*., multiple days) exposures can provide additional insight into cold tolerance, especially if ecologically-relevant conditions are selected (McIntyre et al. 2023; Lemay et al. 2024).

In this study, we characterized the thermal limits of adult *A. tokioensis* collected in late summer from New Brunswick, Canada. We determined their cold tolerance strategy and survival following exposure to acute (short) and chronic mild low temperatures. We also measured CTmin, SCP, and tissue glucose concentrations in worms that were acclimated to summer-like or fall-like conditions. Our results strongly suggest that adult *A. tokioensis* have limited cold tolerance, and that cool fall conditions may limit their spread if adults cannot lay cocoons before they are killed by low temperatures.

## Methods

### Worm collection, identification, and acclimation

Approximately 200 adult jumping worms were collected from a residential back yard near Oromocto, New Brunswick, Canada (45.84°N; 66.48°W) in August 2023 and transported to Saint Francis Xavier University in Antigonish, Nova Scotia, Canada within a few hours. Jumping worms were collected from under logs, plant pots, and other material in the yard. Adults were distinguished from juveniles based on the presence of a clitellum. Recent research found only *A. tokioensis* at this location (Bennett et al. 2024), but we also looked for spermathecal pores on some of the collected earthworms to confirm that they were *A. tokioensis* (Chang et al. 2016). Worms were subdivided into cylindrical 1 L plastic jars with screw-top lids containing 500 – 700 mL moist potting soil and dried aspen leaves (collected the previous fall in Antigonish) for food. Lids were perforated for aeration, and the jars were placed in fine mesh bags to contain escapees. Soil moisture and leaves were checked and topped up twice per week.

Worms were separated into three groups based on the duration and temperature exposures in lab (Table 1). “Summer-collected” worms were used in experiments within 2 – 5 days of field collection. “Summer-acclimated” worms were kept under summer-like conditions (room temperature, c. 20 °C, and natural lighting) for three weeks prior to experiments. “Fall-acclimated” worms were moved into a MIR-154-PA incubator (PHCbi, Chicago, IL, USA) for a five-week fall-like acclimation in darkness prior to experiments. The fall-like acclimation started with one week at 18 °C, and then decreased once per week to 15, 12, 10, and 8 °C to represent average soil temperatures from late September to late October near the collection site (Environment Canada 2023). The samples sizes of each group and their use in subsequent experiments are summarized in Table 1.

**Table 1.** Samples sizes of *Amynthas tokioensis* used in acclimations and experiments^a^ for this study, including mortality that occurred during acclimations.

| Number of worms | Summer-collected | Summer-acclimated | Fall-acclimated |
| --- | --- | --- | --- |
| Total | 16 | 95 | 91 |
| Died prior to experiments | N/A<br>(0% mortality) | 30<br>(32% mortality) | 71<br>(78% mortality) |
| Used in experiments | 16 (cold tolerance strategy) | 32 (acute cold shock)<br>12 (prolonged chilling)<br>16 (CTmin and SCP)<br>5 (glucose assay) | 8 (acute cold shock)<br>6 (CTmin and SCP)<br>6 (glucose assay) |

<sup>a</sup>CTmin = critical thermal minimum, SCP = supercooling point

### Cold tolerance strategy and low temperature tolerance

To determine the impact of cold exposure on worm survival, we conducted three types of cold exposures. First, we determined whether summer-collected worms could survive freezing and supercooling in a cold tolerance strategy experiment. Second, we measured survival following exposure to a range of acute (1 h) cold shocks in summer- and fall-acclimated worms. Third, we tracked survival of summer-acclimated worms during a prolonged (multi-day) exposure to mild low temperatures.

To determine their cold tolerance strategy, we gradually cooled (at −0.25 °C/min) a group of worms from room temperature (c. 20 °C) to a temperature (c. −4 °C) at which half of the worms froze and half remained unfrozen (supercooled), and assessed survival following methods similar to those used for insects (Sinclair et al. 2015; Li et al. 2020; McIntyre et al. 2023; Lemay et al. 2024). Two groups of eight worms were used, both summer-collected. Freezing was detected based on a sudden increase in worm body temperature due to the exothermic process of ice formation (Sinclair et al. 2015). After the cold treatment, worms were returned to room temperature containers with moist soil and leaves for recovery. Survival was assessed as the ability of worms to move independently or respond to gently prodding with a paintbrush 48 h post-cold treatment. Worms were classified as freeze-tolerant if all worms (frozen and supercooled) survived, or freeze-intolerant if only supercooled worms survived.

To conduct the cold exposure described above, we used a programmable recirculating container with a custom attachment. Prior to the cold exposure, each worm was briefly rinsed with water, weighed, and placed into a 50 mL Falcon tube (VWR, Toronto, ON, Canada) without soil. The lack of soil was to ensure that freezing occurred internally, rather than due to inoculation by external ice (*e*.*g*., in frozen soil; Holmstrup 2003; Toxopeus and Sinclair 2018). These tubes were secured in a custom clear container, through which temperature-controlled 50% (v/v) propylene glycol (VWR) could be circulated. The propylene glycol was cooled with an Arctic 25 programmable recirculating chiller (Thermo Fisher Scientific, Toronto, ON, Canada), and pumped into the custom container via an EcoPlus aquarium pump (1 Fish 2 Fish, Dartmouth, NS, Canada). The temperature in the bottom of each tube was recorded once per second via a Type T copper-constantan thermocouple (Omega Engineering, Norwalk, CT, USA) threaded through a hole in the top of the Falcon tube, and interfaced with PicoLog v. 6.2.8 via a TC-8 unit (Pico Technology, Cambridgeshire, UK).

Survival following acute (1 h) cold shock was assessed at temperatures that encompassed 0 – 100% mortality, followed by return to room temperature for recovery. Four groups of eight summer-acclimated worms were used – one group per temperature (−2, 0, 2, and 5 °C). In addition, one group of eight fall-acclimated worms were exposed to 0 °C. The cold exposure was conducted similarly to that described above, except that the temperature was not gradually decreased; worms were transferred directly from room temperature to the cold temperature, and then back to room temperature. In addition, we included c. 20 mL of moist potting soil in each tube prior to cold exposure to make the environment more realistic. We performed a logistic regression in R v4.0.3 (R Core Team 2023) to describe the relationship between temperature and proportion survival in summer-acclimated worms. This experiment allowed us to determine whether short exposures to temperatures above the SCP caused mortality, helping distinguish between freeze-avoidant (high survival) and chill-susceptible (low survival) classifications.

We also assessed survival of summer-acclimated worms during a longer exposure to 5 °C, a temperature that has previously been reported as limiting for *Amynthas* adults. One group of 12 worms was used, and survival was checked after 1, 3.5, 6, 24, and 48 h of chilling. The cold exposure was conducted using a walk-in 5 °C incubator in complete darkness. Temperature was monitored in the incubator by an iButton (iButton Link Technology, Whitewater, WI, USA). Each worm was placed in a 50 mL Falcon tube containing c. 20 mL of moist soil and at least one dried leaf. Each survival assessment required approximately 15 min at room temperature. We performed a logistic regression in R v4.0.3 to describe the relationship between time at 5 °C and proportion survival. This experiment allowed us to determine whether longer exposures to temperatures about the SCP caused mortality, providing further context for whether the worms were chill-susceptible.

### Critical thermal minimum (CTmin) and supercooling point (SCP)

To determine the impact of acclimation on cold tolerance, we measured CTmin and SCP in both summer-acclimated and fall-acclimated worms using a gradual chilling protocol. Using similar methods as described above, we cooled (at −0.25 °C/min) groups of 6 to 8 worms from 20 °C (if summer-acclimated) or 8 °C (if fall-acclimated) to a temperature (c. −6 °C) at which all the worms froze. Prior to the cold exposure, worms were weighed, rinsed, and placed into tubes that did not contain soil, as we needed to observe worm movement. Each worm was observed for activity once every 3 min, and the CTmin was defined as the highest temperature at which voluntary movement ceased for at least 9 min (three observation periods). Voluntary movement included whole body movement (*e*.*g*., climbing the sides of the tube), and small movements (*e*.*g*., of mouthparts) that indicated some part of the neuromuscular system was functional. Cooling continued below the CTmin until the SCP was reached, *i*.*e*., the lowest temperature prior to the exotherm associated with internal ice formation (Sinclair et al., 2015). Because activity and freezing can be impacted by mass, we used ANCOVAs in R v4.0.3 with mass as a covariate to determine the effect of acclimation on CTmin and SCP.

### Worm homogenization and glucose assays

To investigate a biochemical correlate of cold tolerance, we determined concentrations of the potential cryoprotectant glucose in homogenates of whole worms that were summer-acclimated or fall-acclimated, but never exposed to a cold treatment. First, five summer-acclimated worms and six fall-acclimated worms were rinsed with water, flash-frozen in liquid nitrogen in 15 mL

Falcon tubes, and stored for up to three weeks at −80 °C. Worms were then partially thawed on ice, dissected in a petri plate to remove gut contents (which appeared to inhibit the glucose assay in a pilot experiment), and cut into pieces c. 1 cm long. We reweighed the tissues in a 15 mL Falcon tube (one per worm), and added 10 mL of Tris-buffered saline (TBS; 5 mM Tris; 137 mM NaCl; 2.7 mM KCl, pH 6.6) per gram of worm tissue. We used a Power Gen 125 Homogenizer (Thermo Fisher Scientific) two to three times for 10 s at 18000 rpm to homogenize the tissues of each worm. To create a cell-free extract, we centrifuged the homogenate for 5 min at 3000 × *g* at 4 °C, and then centrifuged 1 mL of the resulting supernatant for 30 min at 3000 × *g* at 4 °C.

We performed glucose assays using the Glucose Assay Reagent (Sigma Aldrich, Toronto, ON, Canada) according to the manufacturer’s instructions and as described previously (Toxopeus et al. 2019a). Briefly, we prepared glucose standards in TBS ranging from 0.01 to 0.16 mg/mL via a 2-fold dilution series. We diluted cell-free homogenates 1:1 with TBS, and then incubated these samples (and the standards) at 70 °C for 10 min to denature any endogenous enzymes, followed by centrifugation for 3 min at 20,000 × *g* at room temperature to pellet any precipitated proteins. 90 μL Glucose Assay Reagent and 10 μL sample or standard were pipetted in triplicate into a 96-well spectrophotometer plate. The plate was sealed with Parafilm, followed by mixing of the plate for 30 s on low in a SpectraMax iD5 (Molecular Devices), and incubation of the plate at room temperature for 20 min. The Parafilm was then removed, followed by reading absorbance at 340 nm. The concentration of glucose in each sample was determined by comparison to the standard curve. We compared the glucose concentrations of summer- and fall-acclimated worms using a Welch’s t-test in R v4.0.3.

### Data and code availability

All figures were produced in R v4.0.3. All data and code for statistical analysis and figure generation are available at https://github.com/jtoxopeus/jumping-worm-cold-tolerance.

## Results

### Amynthas tokioensis are chill-susceptible

Overall, adult *A. tokioensis* collected during the summer showed poor cold tolerance. We can conclude they were freeze-intolerant because worms gradually cooled to their SCP (i.e., those that froze) died, while supercooled (never frozen) worms exposed to the same temperature survived (Table 2), and indeed were active within minutes of being returned to room temperature. When summer-acclimated worms were exposed to mild low temperatures for durations of 1 h, survival was low (< 50%) at 0 °C and below (Fig. 1A). In addition, survivors from the 0 °C treatment did not regain full motility, but could respond to prodding. However, a short exposure to 2 °C or 5 °C was non-lethal (Fig. 1A), and worms regained full motility. When summer-acclimated worms were exposed to 5 °C for longer durations, survival was high during the first 6 h of exposure, but dropped to almost zero after 24 h, and all worms were dead after 48 h (Fig. 1B). We therefore conclude that adult *A. tokioensis* are chill-susceptible.

**Table 2.** Survival and mass of summer-collected *Amynthas tokioensis* cooled to the same low temperature (c. −4°C) at which half of them froze and half remained unfrozen (supercooled).

| Metric | Frozen | Supercooled |
| --- | --- | --- |
| Number of worms | 8 | 8 |
| Number of worms that survived | 0 | 8 |
| Average mass ± s.e. of worms (g) | $0.93 \pm 0.12$ | $0.86 \pm 0.08$ |

**Fig. 1.**
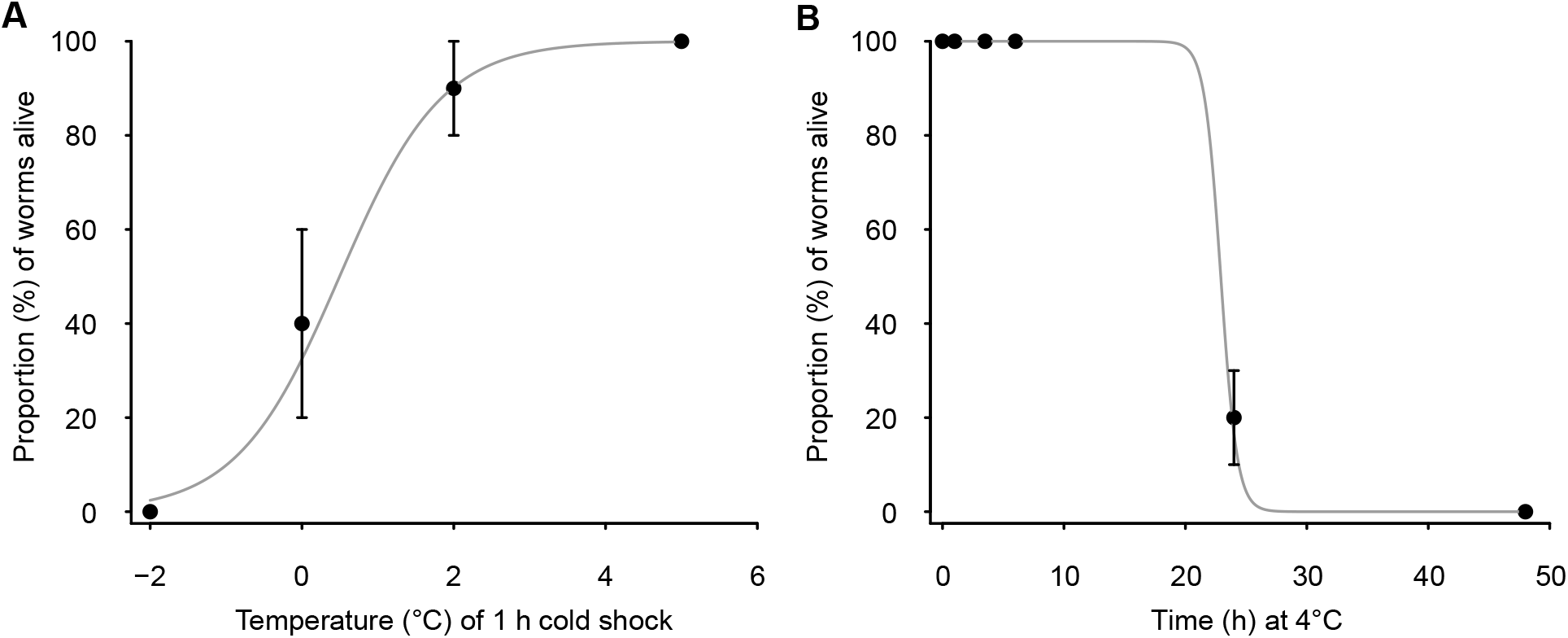
Survival of summer-acclimated *Amynthas tokioensis* following exposure to various cold treatments. **(A)** Each group of 8 worms was cold-shocked for 1 h at one of the indicated temperature. **(B)** One group of 12 worms was kept at 5 °C for up to 48 h, and checked for survival at several time points. Grey lines are based on the logistic regression for each dataset. Temperature significantly impacted survival (regression; *P* = 0.006). Time at 5 °C did not have a statistical impact on survival (regression; *P* > 0.05). Errors bars represent standard error of proportion.

### Amynthas tokioensis do not acclimate well to fall-like conditions

When we exposed *A. tokioensis* to fall-like conditions, the worms did not become more cold-tolerant. Most worms died during the fall-like acclimation (mortality of 78%), while most worms lived during the summer-like acclimation (mortality of 32%; Table 1). Fall-acclimation treatment was associated with a minor decrease in CTmin compared to the summer-acclimation treatment (Fig. 2A; ANCOVA: Mass *P* = 0.92; Acclimation *P* = 0.021). Therefore, fall-acclimated worms (mean CTmin = −0.77 °C) could remain active at temperatures c. 1 °C lower than that of summer-acclimated worms (mean CTmin = 0.63 °C). However, when we exposed six fall-acclimated worms to 0 °C for 1 h, all of them died, exhibiting worse cold tolerance than the summer-collected worms (compare to Fig. 1A). There was no difference in the SCP of fall-acclimated and summer-acclimated worms (mean SCPs = −4.63 °C and −4.37 °C, respectively), although there was an effect of worm mass on SCP (Fig. 2A, ANCOVA: Mass *P* = 0.043; Acclimation *P* = 0.499). Finally, fall-like acclimation didn’t induce an increase in concentration of glucose, a known cryoprotectant in worms (Fig. 2B, Welch’s t-test: *P* = 0.941). Therefore, most of our data support the conclusion that the worms were unable to improve their cold tolerance physiology in response to mild decreases in temperature that would be expected during the fall season.

**Fig. 2.**
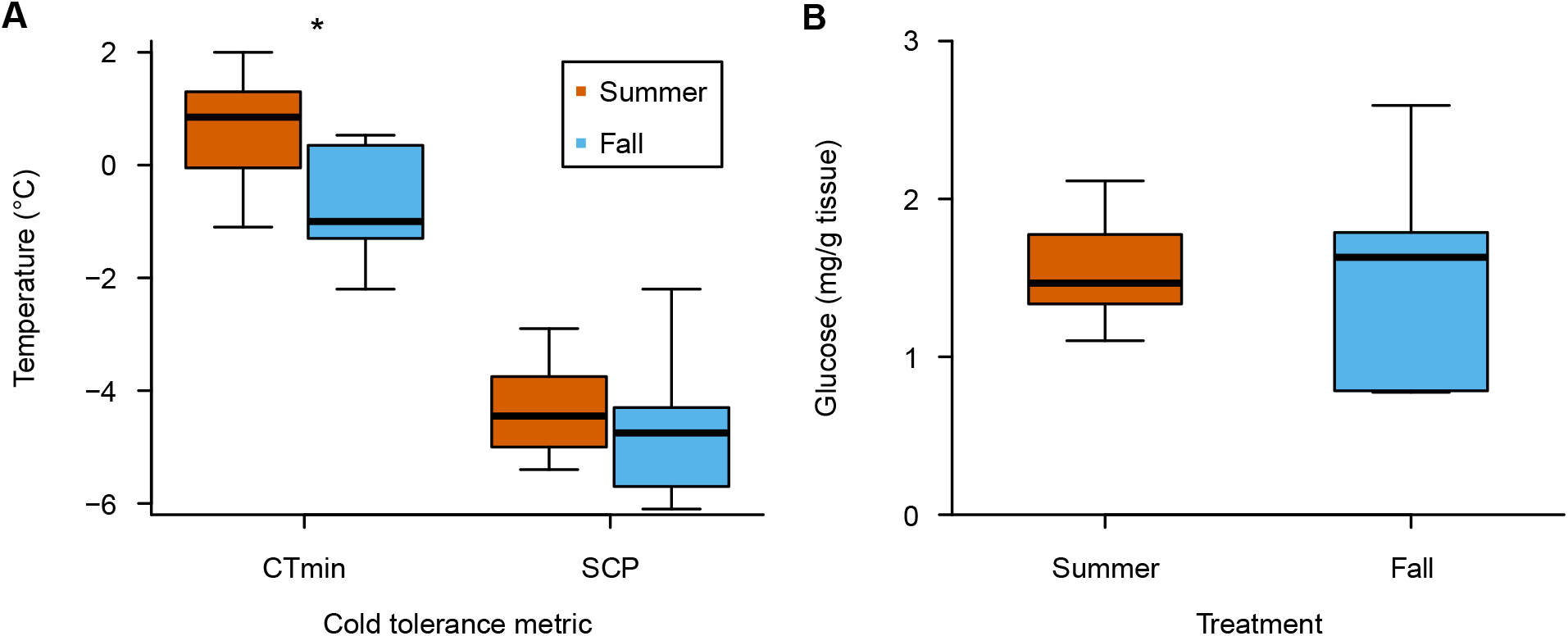
Cold tolerance and cryoprotectant concentrations of summer-acclimated and fall-acclimated *Amynthas tokioensis*. **(A)** A group of 16 (summer) and 6 (fall) worms were gradually chilled to determine the temperature at which voluntary movement ceased (CTmin) and internal ice formation began (SCP). **(B)** Glucose concentration in whole worms (without gut contents) was determined for 5 (summer) and 6 (fall) worms direct from their acclimation conditions. Thick lines represent median, while the bottom and top of the box indicate the first and third quartiles. Whiskers extend to the minimum and maximum y-values. Asterisks indicate a significant difference between summer and fall worms based on ANCOVAs for CTmin and SCP, or Welch’s t-test for glucose concentrations.

## Discussion

Our study is the first to thoroughly characterize the cold tolerance physiology of adult *A. tokioensis*. When gradually cooled, they lost neuromuscular coordination at c. 0 °C, so in their natural environment we would expect these worms to behaviourally avoid temperatures below 0 °C as much as possible (Holmstrup 2003). In the absence of soil, the worms froze at c. −4.5 °C, a relatively high SCP that may be initiated by gut contents – as seen in some insects (Toxopeus et al. 2016, 2019b). In their natural environment these worms could freeze at even higher subzero temperatures if in contact with frozen soil (Holmstrup 2003; Holmstrup et al. 2007). Unlike some earthworms in temperate climates (e.g., *D. octaedra, E. nordenskioldi*; Holmstrup et al. 1999; Rasmussen and Holmstrup 2002), adult *A. tokioensis* did not survive internal ice formation. This is consistent with their relatively tropical origin (Chang et al. 2021), where we would not expect freeze tolerance to evolve (Toxopeus and Sinclair 2018). Based on survival following multiple types of chilling and cold exposures, we conclude that these worms are chill-susceptible.

Despite the poor cold tolerance of adult *A. tokioensis*, their population has been persisting at our collection location in New Brunswick for a few years (McAlpine et al. 2022). Similar to the co-invasive adult *A. agrestis* (Richardson et al. 2009; Görres et al. 2014, 2016), our worms exhibited poor survival at mild low temperatures, showing recovery from short (1 h) chilling at 2 and 5 °C, but not prolonged (1 day or more) exposures. This is consistent with field observations that abundance of adult jumping worms in the field declines as soil temperatures drop below 5 °C (Chang et al. 2021). Similar to many earthworms (Holmstrup 1994; Meshcheryakova and Berman 2014), it is probable that the cocoons of *A. tokioensis* are much more cold-tolerant than the adults. Hatchlings have been observed in early spring in Vermont following a winter in which the soil was frozen, indicating that cocoons can survive through cold winter conditions although it is unclear how winter temperatures impact hatching success rates (Görres et al. 2016). We predict that *A. tokioensis* can persist wherever the growing season is sufficiently warm and long enough for the adults to mature, reproduce, and lay cocoons prior to early fall.

One question that remains is whether the decline in adult *A. tokioensis* abundance each fall is due to accumulation of cold/chilling injuries (Overgaard and MacMillan 2017) or age. Mortality was high during our fall-like acclimation, suggesting early fall (late September to late October) temperatures are stressful. In addition, our worms showed a general inability to modify their physiology (CTmin, SCP, glucose concentrations) during fall-like acclimation, which is characteristic of chill-susceptible earthworms (Holmstrup et al. 1999; Meshcheryakova and Berman 2014). However, the fall-like acclimation was relatively long (5 weeks), so we acknowledge that some worms may have simply died from old age. Jumping worms likely survive as adults for no more than three months (Chang et al. 2021). *Amynthas agrestis* adults can persist under laboratory conditions for up to 4 weeks at moderate temperatures (12 °C; Richardson et al. 2009), and longer acclimations have been successfully used for freeze-tolerant earthworms (e.g., 8 weeks; Holmstrup et al. 2007). A more detailed time course of survival during fall-like conditions would further add to our understanding of how cold impacts the adults of *A. tokioensis*.

## Acknowledgements

The authors would like to thank Samantha Bennett, Carol Dalley, Helen Phillips, and Alyssa Rice for help with field collections of the worms, Steven MacDonald for building the custom clear container that we used for most cold exposures, Luke Burton and Tammy Rodela for optimizing use of the clear container, and Anne Dalziel for connecting EKC and JT. This work was funded by a New Brunswick Wildlife Trust Fund Grant to EKC, and Natural Sciences and Engineering Research Council of Canada (NSERC) Discovery Grants to EKC and JT.

## Statements and Declarations

### Author Contributions

All authors contributed to study conception and design. Data collection was performed by VEA, SRC, TMC, EG, ALG, BEMS, and JT. Data analysis was performed by JT. The first draft of the manuscript was written by JT and EKC, and all authors contributed to pre-submission versions of the manuscript. All authors read and approved the final manuscript.

## References

Bennett S, Phillips HRP, Dalziel AC, et al (2024) Testing the impacts of invasive jumping worms at their northern range limit. Eur J Soil Biol 120:103590. 10.1016/j.ejsobi.2023.103590

Blackmon JH, Taylor MK, Carrera-Martínez R, et al (2019) Temperature affects hatching success of cocoons in the invasive Asian earthworm Amynthas agrestis from the Southern Appalachians. Southeast Nat 18:270–280. 10.1656/058.018.0205

Chang C-H, Bartz MLC, Brown G, et al (2021) The second wave of earthworm invasions in North America: biology, environmental impacts, management and control of invasive jumping worms. Biol Invasions 23:3291–3322. 10.1007/s10530-021-02598-1

Chang C-H, Johnston MR, Görres JH, et al (2018) Co-invasion of three Asian earthworms, Metaphire hilgendorfi, Amynthas agrestis and Amynthas tokioensis in the USA. Biol Invasions 20:843–848. 10.1007/s10530-017-1607-x

Chang C-H, Snyder BA, Szlavecz K (2016) Asian pheretimoid earthworms in North America north of Mexico: An illustrated key to the genera Amynthas, Metaphire, Pithemera, and Polypheretima (Clitellata: Megascolecidae). Zootaxa 4179: 495–529. 10.11646/zootaxa.4179.3.7

Environment Canada (2023) Historical Climate Data. https://climate.weather.gc.ca/index_e.html. Accessed 20 Aug 2023

Görres JH, Bellitürk K, Melnichuk RDS (2016) Temperature and moisture variables affecting the earthworms of genus Amynthas Kinberg, 1867 (Oligachaeta: Megascolecidae) in a hardwood forest in the Champlain Valley, Vermont, USA. Appl Soil Ecol 104:111–115. 10.1016/j.apsoil.2015.10.001

Görres JH, Melnichuk RDS, Bellitürk K (2014) Mortality pattern relative to size variation within Amynthas agrestis (Goto & Hatai 1899) (Oligochaeta: Megascolecidae) population in the Champlain Valley of Vermont, USA. Megadrilogica 16:9–14

Havird JC, Neuwald JL, Shah AA, et al (2020) Distinguishing between active plasticity due to thermal acclimation and passive plasticity due to Q10 effects: Why methodology matters. Funct Ecol 34:1015–1028. 10.1111/1365-2435.13534

Holmstrup M (1994) Physiology of cold hardiness in cocoons of five earthworm taxa (Lumbricidae: Oligochaeta). J Comp Physiol B 164:222–228. 10.1007/BF00354083

Holmstrup M (2003) Overwintering adaptations in earthworms. Pedobiologia 47:504–510. 10.1078/0031-4056-00220

Holmstrup M, Costanzo JP, Lee RE (1999) Cryoprotective and osmotic responses to cold acclimation and freezing in freeze-tolerant and freeze-intolerant earthworms. J Comp Physiol B 169:207–214. 10.1007/s003600050213

Holmstrup M, Overgaard J, Bindesbøl A-M, et al (2007) Adaptations to overwintering in the earthworm Dendrobaena octaedra: Genetic differences in glucose mobilisation and freeze tolerance. Soil Biol Biochem 39:2640–2650. 10.1016/j.soilbio.2007.05.018

Holmstrup M, Westh P (1994) Dehydration of earthworm cocoons exposed to cold: a novel cold hardiness mechanism. J Comp Physiol B 164:312–315. 10.1007/BF00346448

Holmstrup M, Zachariassen KE (1996) Physiology of cold hardiness in earthworms. Comp Biochem Physiol A 115:91–101. 10.1016/0300-9629(96)00010-2

Lemay K, Moore M, Brown P, et al (2024) Conserved cold tolerance of Rhagoletis species from different host fruits, elevations in Colorado, USA. Physiol Entomol. 10.1111/phen.12439

Li NG, Toxopeus J, Moos M, et al (2020) A comparison of low temperature biology of Pieris rapae from Ontario, Canada, and Yakutia, Far Eastern Russia. Comp Biochem Physiol A 242:110649. 10.1016/j.cbpa.2020.110649

McAlpine D, Reynolds J, Manzer L, Elton K (2022) First reports of invasive pheretimoid earthworms (Oligochaeta: Megascolecidae) of Asian origin in Atlantic Canada. BioInvasions Rec 11:830–838. 10.3391/bir.2022.11.4.02

McIntyre T, Andaloori L, Hood GR, et al (2023) Cold tolerance and diapause within and across trophic levels: Endoparasitic wasps and their fly host have similar phenotypes. J Insect Physiol 146:104501. 10.1016/j.jinsphys.2023.104501

Meshcheryakova EN, Berman DI (2014) Cold hardiness and geographic distribution of earthworms (Oligochaeta, Lumbricidae, Moniligastridae). Entomol Rev 94:486–497. 10.1134/S0013873814040046

Moore J-D, Reynolds J (2024) First record of the invasive Asian earthworm Amynthas tokioensis (Beddard, 1892) in the province of Québec, Canada. BioInvasions Rec 13:1–8. 10.3391/bir.2024.13.1.01

Nuutinen V, Butt KR (2009) Worms from the cold: Lumbricid life stages in boreal clay during frost. Soil Biol Biochem 41:1580–1582. 10.1016/j.soilbio.2009.04.019

Overgaard J, MacMillan HA (2017) The integrative physiology of insect chill tolerance. Annu Rev Physiol 79:187–208

R Core Team (2023) R: A Language and Environment for Statistical Computing. Vienna, Austria: R Foundation for Statistical Computing. Available from http://www.R-project.org

Rasmussen L, Holmstrup M (2002) Geographic variation of freeze-tolerance in the earthworm Dendrobaena octaedra. J Comp Physiol B 172:691–698. 10.1007/s00360-002-0298-4

Reynolds J (2014) A checklist by counties of earthworms (Oligochaeta: Lumbricidae, Megascolecidae and Sparganophilidae) in Ontario, Canada. Megadrilogica 16:111–135

Reynolds JW (2018) First earthworm (Annelida: Oligochaeta) species’ collections in Canada and the continental United States. Megadrilogica 23:1–50

Richardson DR, Snyder BA, Hendrix PF (2009) Soil moisture and temperature: Tolerances and optima for a non-native erthworm species, Amynthas agrestis (Oligochaeta: Opisthopora: Megascolecidae). Southeast Nat 8:325–334. 10.1656/058.008.0211

Sinclair BJ, Coello Alvarado LE, Ferguson LV (2015) An invitation to measure insect cold tolerance: Methods, approaches, and workflow. J Therm Biol 53:180–197. 10.1016/j.jtherbio.2015.11.003

Slotsbo S, Maraldo K, Malmendal A, et al (2008) Freeze tolerance and accumulation of cryoprotectants in the enchytraeid Enchytraeus albidus (Oligochaeta) from Greenland and Europe. Cryobiology 57:286–291. 10.1016/j.cryobiol.2008.09.010

Toxopeus J, Koštál V, Sinclair BJ (2019a) Evidence for non-colligative function of small cryoprotectants in a freeze-tolerant insect. Proc Royal Soc B 286:20190050. 10.1098/rspb.2019.0050

Toxopeus J, Lebenzon JE, McKinnon AH, Sinclair BJ (2016) Freeze tolerance of Cyphoderris monstrosa (Orthoptera: Prophalangopsidae). Can Entomol 148:668–672. 10.4039/tce.2016.21

Toxopeus J, McKinnon AH, Štětina T, et al (2019b) Laboratory acclimation to autumn-like conditions induces freeze tolerance in the spring field cricket Gryllus veletis (Orthoptera: Gryllidae). J Insect Physiol 113:9–16. 10.1016/j.jinsphys.2018.12.007

Toxopeus J, Sinclair BJ (2018) Mechanisms underlying insect freeze tolerance. Biol Rev 93:1891–1914. 10.1111/brv.12425

